# Topoisomerase I inhibition suppresses nuclear blebbing via RNA Pol II stalling and nuclear stiffening

**DOI:** 10.64898/2026.01.09.698681

**Authors:** Nickolas Borowski, Andy Li, Andrew D. Stephens

## Abstract

Abnormal nuclear blebbing occurs in many human diseases and causes nuclear rupture and dysfunction. Nuclear blebbing is caused by chromatin motion via RNA Pol II transcriptional activity and nuclear mechanical weakening. Camptothecin, a topoisomerase I inhibitor, rapidly suppresses nuclear blebbing within hours. We find that camptothecin does not decrease RNA Pol II phosphorylation, but does decrease newly synthesized RNA, likely by stalling RNA Pol II. However, camptothecin treatment suppresses nuclear blebbing more drastically than inhibition of transcription activity by alpha amanitin, suggesting a second mechanism of nuclear blebbing suppression. Dual micromanipulation nuclear force measures revealed camptothecin treatment increased chromatin-based nuclear stiffness but not lamin-based strain stiffening. Thus, inhibition of topoisomerase I via camptothecin drastically suppresses nuclear blebbing by both stalling RNA Pol II and increasing chromatin-based nuclear stiffness.

**Summary statement:** Inhibition of topoisomerase I suppresses nuclear blebbing by stalling RNA Pol II activity and increasing chromatin-based nuclear spring constant.

## Introduction

The nucleus must maintain its shape and integrity to ensure proper function. Abnormal nuclear morphology is a hallmark of many human diseases (Gisselsson et al., 2001; Kalukula et al., 2022). One type of abnormal nuclear morphology is a nuclear bleb, a herniation of the nucleus >1 μm in size with decreased DNA density (Bunner *et al*., 2024; Chu *et al*., 2025; Pujadas Liwag *et al*., 2025). Nuclear blebs lead to nuclear ruptures, and both cause nuclear dysfunction via altered transcription, DNA damage and repair, and cell cycle control (De Vos *et al*., 2011; Helfand *et al*., 2012; Denais *et al*., 2016; Irianto *et al*., 2016; Raab *et al*., 2016; Chen *et al*., 2018; Pfeifer *et al*., 2018; Xia *et al*., 2018; Stephens *et al*., 2019b; Earle *et al*., 2020; Nader *et al*., 2021; Shah *et al*., 2021; Pho *et al*., 2023). Nuclear blebs are the result of actin antagonism deforming a weaker nucleus (Pho et al., 2023; Stephens et al., 2018b) and transcription driven chromatin motion (Berg *et al*., 2023; Prince *et al*., 2025). Nuclear rigidity is determined by the two main mechanical components, chromatin and lamins, and two sub-components, chromatin-chromatin and chromatin-lamin linkers (Banigan et al., 2017; Belaghzal et al., 2021; Hobson et al., 2020; Kalukula et al., 2022; Manning et al., 2025; Nava et al., 2020; Strom et al., 2021; Swift et al., 2013; Williams et al., 2024). The growing understanding of the causes and consequences behind nuclear blebbing prompts further exploration into commonly used cancer drugs that might impact nuclear morphology.

Camptothecin is a well-known inhibitor of topoisomerase I and a widely used anti-cancer drug (Khaiwa et al., 2021; Pommier, 2006). Topoisomerase I is necessary for relieving DNA supercoiling during replication and transcription by generating a single strand break then re-ligation after relaxation (Champoux, 2001). Camptothecin’s topoisomerase I inhibition is documented to inhibit the re-ligation step by causing nicked single strands to remain, leading to replication conflicts and cell death (D’Arpa et al., 1990). However, the effect of camptothecin on nuclear blebbing remains unknown as a possible secondary contributor to its anti-cancer activities. Camptothecin has been reported to disrupt transcription (Ljungman and Hanawalt, 1996; Seiler et al., 2007). Topoisomerase I is a well-known chromatin protein that could have implications in chromatin-based nuclear stiffness. Thus, we aimed to determine the effect of topoisomerase I inhibition by camptothecin on nuclear blebbing, a well-known hallmark of many cancers (Helfand *et al*., 2012; Stephens *et al*., 2018a).

## Results and Discussion

### Topoisomerase I inhibition suppresses nuclear blebbing

Nuclear blebs are defined as >1 µm herniations of the nucleus that have decreased DNA density and occur in interphase (**Fig. 1, A**). First, we imaged HCT116 nuclei for nuclear blebbing percentage in untreated wild type (UNT) at 2.0 ± 0.3%. Chromatin decompaction via overnight treatment with histone deacetylase inhibitor valproic acid (VPA) increased nuclear blebbing to 6.4 ± 0.5% (**Fig. 1B**). Treatment with topoisomerase I inhibitor camptothecin (CPT) for 1 hour suppressed increased nuclear blebbing in VPA treated cells but not in wild type (**Fig. 1B**). Thus, topoisomerase I inhibition via camptothecin rapidly suppressed nuclear blebbing.

**Figure 1.**
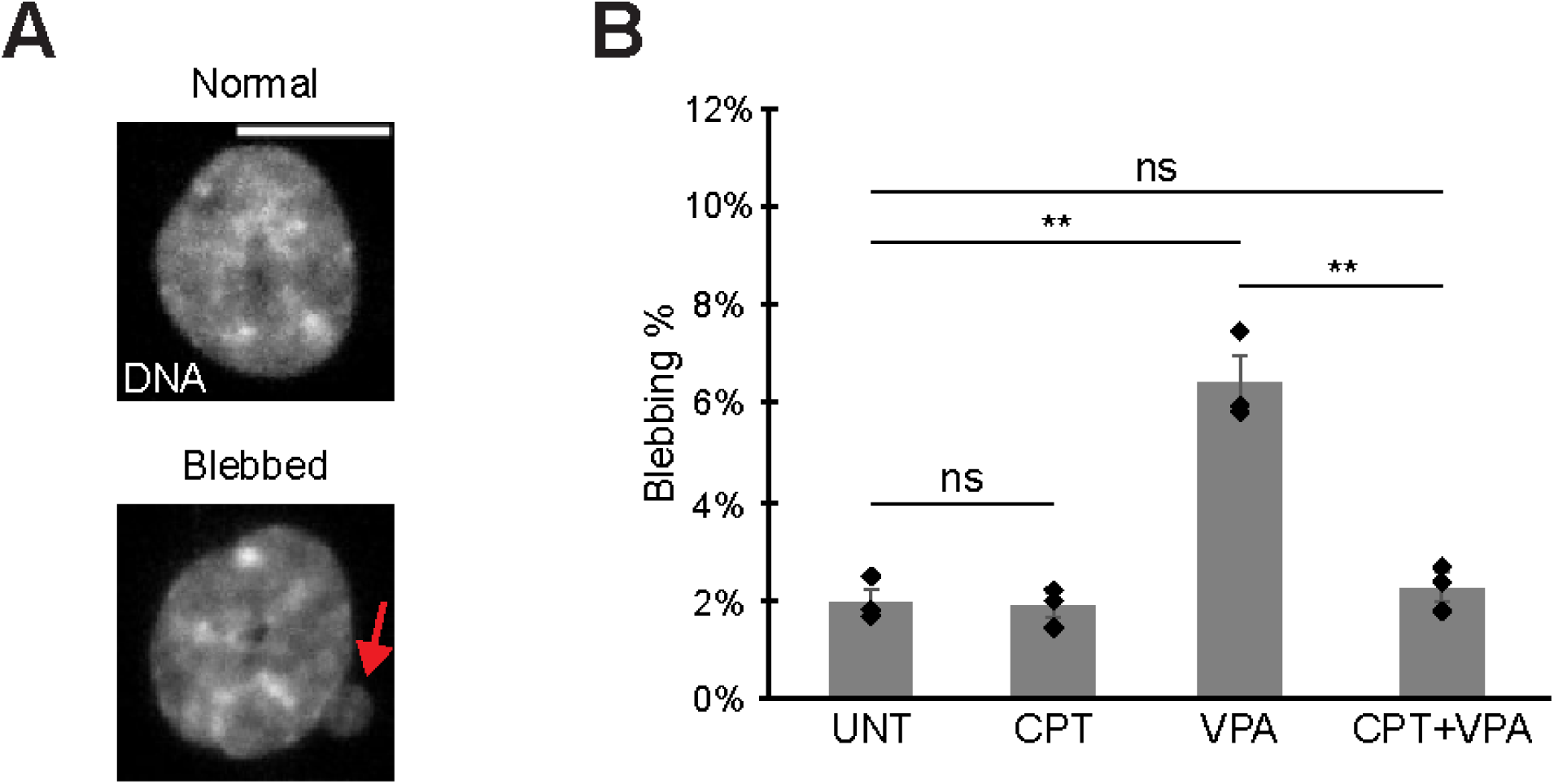
Topoisomerase I inhibition decreases nuclear blebbing. (A) Example images of normal and blebbed HCT116 nuclei with DNA stain Hoechst. (B) Graph of blebbing percentages in HCT116 for wild type untreated (UNT), chromatin decompaction (VPA), camptothecin treated (CPT), and both VPA + CPT. Means of biological triplicates with n > 350 each are shown. The red arrow denotes a nuclear bleb in the example image. Error bars represent standard error. Two-tailed unpaired Student’s t-test between all conditions. P values reported as \**P*<0.05; \*\**P*<0.01; \*\*\**P*<0.001, or ns denotes no significance. Scale bar = 10 µm.

Discussion: Topoisomerase I inhibition by camptothecin is well documented for anti-cancer activities, but our new data uncovers a novel function of camptothecin to suppress nuclear blebbing. Camptothecin’s main functions are to disrupt dividing cells during S-phase DNA replication, causing apoptosis from DNA replication problems and DNA damage (D’Arpa et al., 1990). We find that camptothecin provides a unique secondary anti-cancer activity in which asynchronous cell nuclear blebbing can be suppressed rapidly and drastically, a behavior well-documented to cause dysfunction (Kalukula et al., 2022; Stephens, 2020). Previous studies have reported that camptothecin can disrupt transcription (Bendixen et al., 1990). We have recently reported that inhibition of transcriptional activity can suppress nuclear blebbing (Berg *et al*., 2023; Prince *et al*., 2025). Thus, the data suggests that transcription inhibition is the mechanism causing nuclear bleb suppression in camptothecin-based topoisomerase I inhibition.

### Topoisomerase I inhibition stalls RNA Pol II

Nuclear blebbing relies on RNA Pol II motor activity that can be measured via changes in RNA Pol II activity via immunofluorescence (**Fig. 2, A and B**). Activity of RNA Pol II can be measured via levels of phosphorylation of Ser5 for initiation (Komarnitsky *et al*., 2000; Schwer and Shuman, 2011) and Ser2 for elongation (Barilla *et al*., 2001; Kim *et al*., 2004). Camptothecin treatment did not alter overall RNA Pol II levels as there was no significant change in CRISPR labeled mAID-Clover-POLR2A HCT116 cells (**Fig. 2, A and C**). Transcription initiation-active RNA Pol II pSer5 levels also remained unchanged. Interestingly, RNA Pol II pSer2 elongation levels significantly increased relative to wild type measurements (**Fig. 2, A and C**), suggesting RNA Pol II was stalling in the elongation stage. This data agrees with previous studies finding stalling in elongation by camptothecin (Ljungman and Hanawalt, 1996; Seiler et al., 2007). Overall, camptothecin does not decrease RNA Pol II levels or phosphorylation activity state but does increase RNA Pol II in the elongation state.

**Figure 2.**
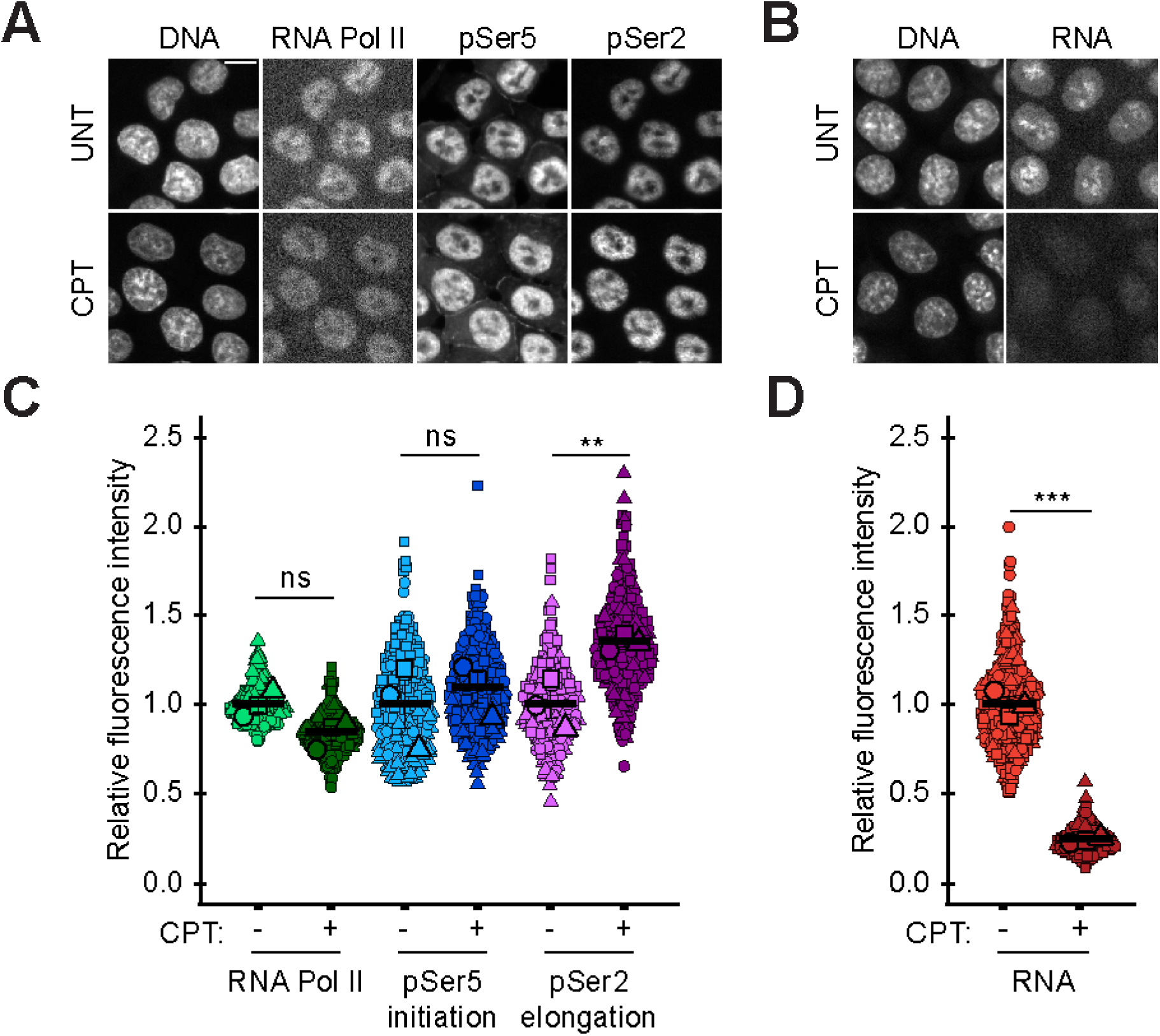
Topoisomerase I inhibition increases RNA Pol II pSer2 and decreases RNA levels. (A-B) Example images and (C-D) graphs of RNA Pol II, RNA Pol II phosphorylated at Ser5 (pSer5, initiation) or Ser2 (pSer2, elongation), and RNA levels via Click-iT chemistry in wild type (-) and camptothecin treated (CPT, +) HCT116 cells. RNA Pol II, pSer5, and pSer2 biological triplicates with n = 30 each, total 90. RNA levels via Click-iT chemistry biological triplicates with n = 40 each, total 120. Statistical tests are one-way ANOVA with a post-hoc Tukey test. For all panels error bars represent standard error, p values reported as \**P*<0.05; \*\**P*<0.01; \*\*\**P*<0.001, or ns denotes no significance. Scale bar = 10 µm.

We hypothesized that topoisomerase I inhibition induced RNA Pol II stalling. To determine RNA Pol II activity output, we measured newly synthesized RNA via Click-iT chemistry. Newly synthesized RNA levels drastically decreased upon camptothecin treatment (**Fig. 2, B and D**). Thus, camptothecin does not decrease RNA Pol II phosphorylation, but does stall RNA Pol II in elongation and hinders RNA production.

Discussion: Transcriptional activity has been shown to be necessary for nuclear blebbing, especially at rates above wild type levels (Berg *et al*., 2023; Prince *et al*., 2025). Previous studies disrupted RNA Pol II activity via decreased levels of RNA Pol II and/or decreased levels of RNA Pol II pSer5/2. Our data showing increased levels of RNA Pol II pSer2 and a drastic decrease in RNA levels suggests that camptothecin stalls RNA Pol II during elongation and RNA production. This was done without decreasing RNA Pol II total or phosphorylated levels. Thus, we reveal a novel method of suppressing RNA Pol II motor activity, which was previously shown to be an important driver of chromatin motion to form and stabilize nuclear blebs.

### Topoisomerase I inhibition suppresses nuclear blebbing more than inhibition of transcriptional activity

We hypothesized that suppression of nuclear blebbing by camptothecin-based topoisomerase I inhibition is due to inhibition of transcriptional activity. To test this hypothesis, we first measured nuclear blebbing in MEF cell lines treated with RNA Pol II inhibitor alpha amanitin or camptothecin. Compared to HCT116 nuclei, MEF nuclei have a higher baseline wild type nuclear blebbing percentage at 3.6 ± 0.5%, allowing us to better determine whether nuclear blebbing can be suppressed in untreated wild type (UNT, **Fig. 3, A and B**). In agreement with numerous publications, VPA treatment causes an significant increase in nuclear blebbing to 8.4 ± 1.1%. Treatment of cells with transcriptional activity inhibitor alpha amanitin (AAM) overnight did not suppress nuclear blebbing in wild type. Alpha amanitin did suppress the increased level of nuclear blebbing upon chromatin decompaction with VPA treatment back to wild type levels, around 4% (**Fig. 3B**). Overall, this data is in strong agreement with our previous publications showing that transcription inhibition can rescue increased nuclear blebbing back to wild type levels (Berg *et al*., 2023; Prince *et al*., 2025).

**Figure 3.**
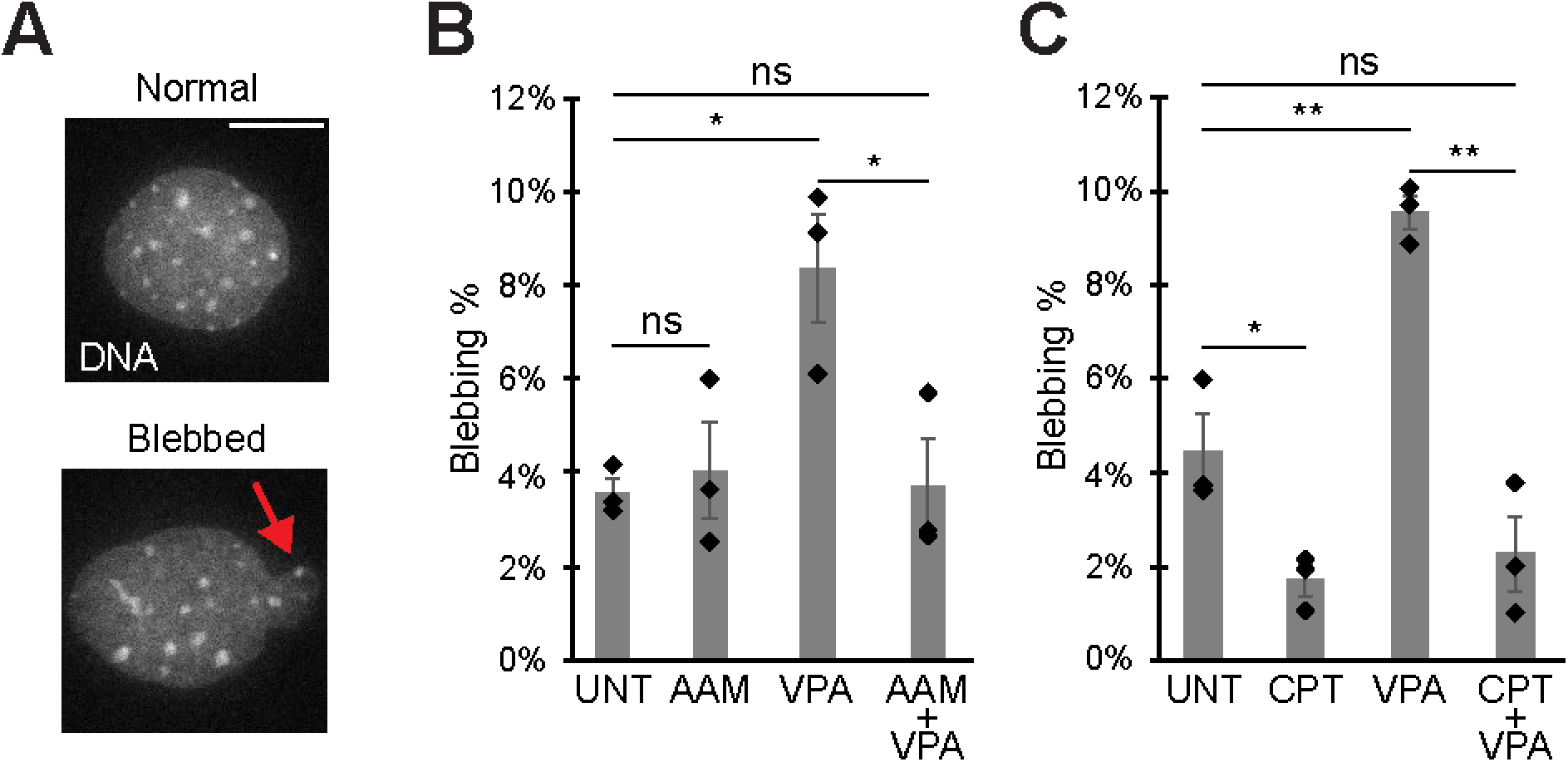
Camptothecin topoisomerase I inhibition suppresses nuclear blebbing more than transcription activity inhibition via alpha amanitin. (A) Example images of normal and blebbed MEF nuclei and (B-C) graphs of blebbing percentages in MEF cells for untreated (UNT), chromatin decompaction (VPA), alpha amanitin treated (AAM), camptothecin treated (CPT), and both VPA + AAM or CPT. Means of biological triplicates with n > 100 total each are shown. The red arrow denotes a nuclear bleb in the example image. Error bars represent standard error. Two-tailed unpaired Student’s t-test between all conditions. P values reported as \**P*<0.05; \*\**P*<0.01; \*\*\**P*<0.001, or ns denotes no significance. Scale bar = 10 µm.

**Figure 4.**
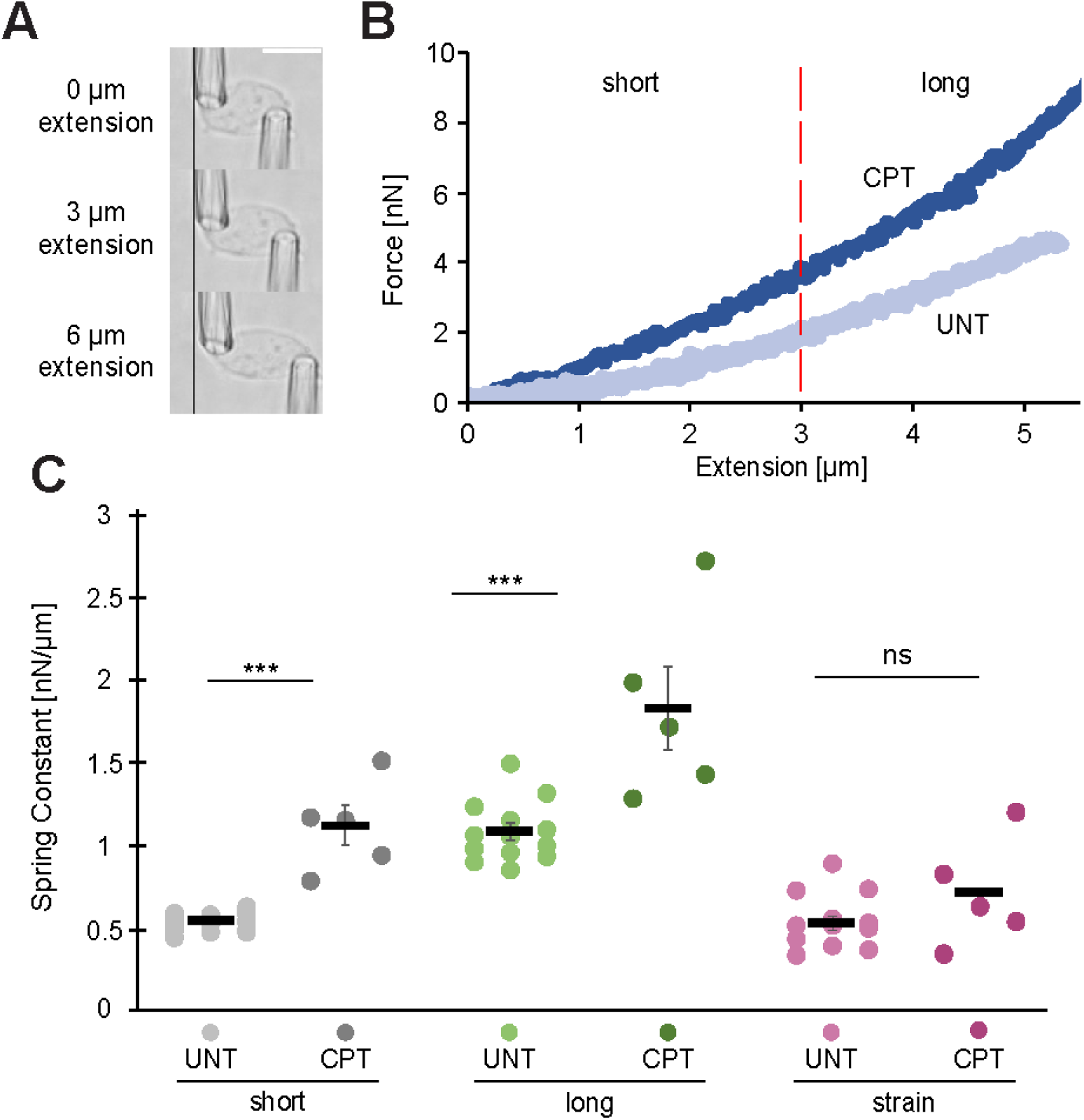
Topoisomerase I inhibition increases nuclear stiffening in MEF. (A) Example images of MEF *V−/−* isolated nucleus dual pipette micromanipulation force measurements. (B) Example graph of force/extension including example wild type and camptothecin treated (CPT). The dashed line denotes the switch from the short-extension regime to the long-extension regime. (C) Graph of individual (dots) and average (black bar) short-extension (‘UNT’, n=18; ‘CPT’, n=5), long-extension (‘UNT’, n=13; ‘CPT’, n=5), and strain-stiffening (‘UNT’, n=13; ‘CPT’, n=5), nuclear spring constants in camptothecin treated and untreated conditions. Two-tailed unpaired Student’s t-test was used. For all panels error bars represent standard error, p values reported as \**P*<0.05; \*\**P*<0.01; \*\*\**P*<0.001, or ns denotes no significance. Scale bar = 10 µm.

Next, we aimed to determine if suppression of nuclear blebbing due to camptothecin-based topoisomerase I inhibition is similar to transcriptional inhibition via alpha amanitin. Similar to the previous experiment, MEF untreated wild type nuclear blebbing percentage was 3.7 ± 0.5% and increases to 10.0 ± 1.0% nuclear blebbing upon VPA treatment (**Fig. 3C**). Camptothecin treatment (CPT) for 4 hours significantly decreased nuclear blebbing in both wild type to 1.7 ± 0.4% and chromatin decompaction via VPA to 2.2 ± 0.8% (**Fig. 3C**). This statistically significant nuclear bleb suppression exceeds that caused by RNA Pol II inhibitor alpha amanitin (**Fig. 3B vs. 3C**). Thus, the topoisomerase I inhibitor camptothecin causes suppression of nuclear blebbing to a greater degree than inhibition of transcriptional activity via alpha amanitin.

Discussion: Camptothecin provides an enhanced level of nuclear blebbing suppression. Previous modulations, including either increased nuclear rigidity (Stephens et al., 2018b; Stephens et al., 2019b; Strom et al., 2021) or decreased transcription (Berg *et al*., 2023; Prince *et al*., 2025), have only been reported to rescue nuclear blebbing back to wild type levels from perturbations that increase nuclear blebbing, like chromatin decompaction and loss of lamins. However, camptothecin-based topoisomerase I inhibition provides greater suppression of nuclear blebbing in both wild type and VPA chromatin decompaction than previously reported. The lack of nuclear bleb suppression in wild type HCT116 cells is likely due to the already low (∼2%) level and shorter treatment time of 1 hour. In comparison, MEF higher baseline wild type nuclear blebbing (∼4%) provides a greater range over which to determine suppression along with longer 4 hour treatment. This strongly suggests that camptothecin is modulating more than a single pathway to suppress nuclear blebbing.

### Topoisomerase I inhibition increases chromatin-based but not lamin-based nuclear spring constant

Another major factor in nuclear blebbing is nuclear rigidity (Lammerding *et al*., 2005; Furusawa *et al*., 2015; Stephens *et al*., 2018b, 2019a; Pho *et al*., 2024). We hypothesized that camptothecin’s greater suppression of nuclear blebbing could be due to changes in nuclear spring constant. We used dual micropipette manipulation of isolated nuclei to measure the nuclear spring constant (**Fig. 3A**). We used MEF vimentin null (*V−/−*) cells for ease of nucleus isolation. MEF *V−/−* nuclei have been shown to have the same nuclear spring constant and nuclear blebbing percentages as wild type (Berg et al., 2023; Pho et al., 2023; Stephens et al., 2017; Stephens et al., 2018b). Treatment with camptothecin resulted in an increased nuclear spring constant (**Fig. 3B**).

Dual micropipette micromanipulation excels at determining changes in chromatin vs. lamin contributions. We find that camptothecin significantly increased both the short extension (chromatin-based) and long extension (chromatin + lamins) nuclear spring constants (**Fig. 3C**). However, lamin-based nuclear strain stiffening did not significantly change, which is determined by subtracting the short from the long regime nuclear spring constant (**Fig. 3C**). Thus, camptothecin provides increased chromatin-based nuclear spring constant that leads to a greater level of nuclear blebbing suppression alongside disruption of RNA Pol II motor activity.

Discussion: Our data strongly supports that inhibition of topoisomerase I causes increased chromatin-based stiffness as the second mechanism of nuclear bleb suppression. Specifically, modulations of transcriptional activity show no change in nuclear spring constant (Berg *et al*., 2023; Prince *et al*., 2025). Thus, in addition to transcription disruption, camptothecin also increased chromatin-based nuclear spring constant. Camptothecin is reported to inhibit the re-ligation of the cut single stranded DNA while topoisomerase I is bound to DNA. This conformation could coat the DNA to strengthen single strands similar to the reported chromatin-based nuclear rigidity that occurs upon addition of DNA binders like Hoechst and propidium iodide (Stephens et al., 2017). Alternatively, this inhibition could introduce chromatin-chromatin crosslinking interactions, which also result in increased chromatin-based nuclear spring constant (Belaghzal et al., 2021; Strom et al., 2021; Williams et al., 2024). While the mechanism for increasing chromatin-based nuclear rigidity remains unclear, this finding provides novel data for follow up investigations. Furthermore, the data solidify that camptothecin ultimately can protect the nucleus through two mechanisms, transcription activity disruption and nuclear rigidity, to suppress nuclear blebbing and its associated dysfunctions.

Conclusion: We provide novel data for secondary functions of the anti-cancer drug camptothecin, a topoisomerase I inhibitor via rapid and drastic suppression of nuclear blebbing. The mechanism of this action involves both stalling of RNA Pol II and increased chromatin-based nuclear spring constant. This finding supports previously established determinants of nuclear blebbing. Furthermore, this highlights the potential for other chromatin drugs to suppress nuclear blebbing, a well-known hallmark and contributor to many human diseases.

## MATERIALS AND METHODS

### Cell growth

HCT116 RNA Pol II AID2 cells (Yesbolatova *et al*., 2020) from the Kanemaki lab (Department of Chromosome Science, National Institute of Genetics, Research Organization of Information and Systems (ROIS), Yata 1111, Mishima, Shizuoka 411-8540 Japan) were cultured in McCoys 5A media (Gibco) complete with Pen Strep (Fisher) and 10% fetal bovine serum (FBS; HyClone) at 37°C and 5% CO2. Upon reaching confluency, cells were trypsinized, replated, and diluted into fresh media every other day. Immortalized Mouse Embryonic Fibroblast (MEF), previously described in (Shimi et al., 2008; Stephens et al., 2018; Vahabikashi et al., 2022), were handled the same but cultured in DMEM (Corning).

### Cell treatments

Cells were plated into eight-well dishes for 48 h prior to imaging. Cells were either left untreated (UNT), treated with valproic acid (VPA (Sigma-Aldrich 1069-66-5)) at 2 mM for 24 h, treated with a transcription altering drug, or dual treated with VPA and a transcription altering drug. Cells were treated with camptothecin (CPT (Thermo Scientific 7689-03-4)) at 10 μM for 1 or 4 hours to achieve topoisomerase I inhibition. Cells were treated with alpha-amanitin (AAM Cayman Chemicals 23109-05-9)) at 10 μM for 24 hours to achieve RNA Pol II inhibition. Cells were treated with Hoechst 33342 at a dilution of 1:10,000 for 15 min before population imaging.

### Immunofluorescence

Cells were plated into eight-well dishes for 48 h prior to fixation. A 1:3 solution of cells and media respectively was made and 300 μL of solution was plated into each well. Cells were treated as mentioned above. Fixation was done with 4% paraformaldehyde (Electron Microscopy Sciences) in PBS for 15 min at room temperature. Cells were then washed 3 times for 10 min each with PBS, permeabilized with 0.1% Triton X-100 (US Biological) in PBS for 15 min, and washed with 0.06% Tween-20 (US Biological) in PBS for 5 min followed by 2 more washes in PBS for 5 min each at room temperature. After that, cells were blocked for 1h at room temperature using a blocking solution consisting of 2% bovine serum albumin (BSA (Fisher Scientific)) in PBS. Primary antibodies were diluted in the blocking solution as follows: anti-RNA pol II CTD repeat YSPTSPS (phospho S5/pSer5) at 1:1000 (ab5408; Abcam), anti-RNA pol II CTD repeat YSPTSPS (phospho S2/pSer2) at 1:1000 (ab5095; Abcam). Cells were incubated in the primary antibodies overnight at 4℃ in the dark and then washed with PBS three times for 5 min each. Cells were then incubated with anti-mouse and anti-rabbit Alexa Fluor 555 and 647 fluorescent secondary antibodies diluted at 1:1000 in blocking solution for 1 h at room temperature in the dark. Cells were washed with Hoechst 33342 (Life Technologies) in PBS at a 1:1000 dilution for 5 min and then washed three more times with PBS. Cells were mounted using Prolong Gold anti-fade mountant (Invitrogen, P36930) and cured overnight in the dark.

### RNA labeling

RNA labeling was accomplished using Click-iT RNA Alexa Fluor 594 Imaging Kit (Invitrogen, C10330). Cells were fixed using the same immunofluorescence protocol listed above, with the following modifications for RNA labeling: prior to cell fixation, cells were incubated for 1 h using 5-ethynyl uridine (EU) at a final concentration of 1 mM. Cells were then washed three times with PBS, fixed, and then permeabilized. After permeabilization, 300 μL of the Click-iT reaction cocktail was added and incubated for 30 mins in the dark. The reaction cocktail was then removed and cells were washed with the Click-iT reaction rinse buffer for 5 min. The reaction buffer was then removed and cells were washed 3 times with PBS, 5 min each. Cells were then stained with Hoechst 33342 (Life Technologies) in PBS at a 1:1000 dilution for 5 min and then washed three more times with PBS. Cells were kept and imaged in PBS or mounted using Prolong Gold anti-fade mountant (Invitrogen, P36930) and left to cure overnight in the dark.

### Michromanipulation force measurement of an isolated nucleus

Micromanipulation force measurements were conducted as described in (Currey et al., 2022; Stephens et al., 2017). Vimentin null (*V*−/−) MEF cells were used for their ease of nucleus isolation and have similar nuclear rigidity to wild-type nuclei. MEF *V*−/− cells were grown in micromanipulation wells to provide micropipettes with low angle access to the isolated nuclei. Cells were either left untreated or were treated with CPT at 10 µm for 4 h. A Flaming/Brown micropipette puller p-97 (Sutter Instruments) is used to create micropipette. The pull and spray pipettes are made using World Precision Instruments TW100-6 and force pipettes are made using TW100F-6. The pull program used to create the pull and spray pipettes is as follows: Heat 564, Pull 110, Velocity 110, Time 100, and Pressure 500. The pull program used to create the force pipettes is as follows: Heat 561, Pull 220, Velocity 200, Time 20, and Pressure 500. Pulling completes with the separation of the capillary into two pipettes, which are cut with a heated filament to an ideal width of 3-6 µm.

We used our previously published micromanipulation set up consisting of Nikon Ts2R-FL, 10X and 60 phase objectives, and Amscope MU130 camera along with two Sutter Instruments MP-285 (Currey et al., 2022). MEF *V*−/− cell nuclei were isolated via spray micropipette of the detergent Triton X-100 (0.05%) in PBS and the “pull” micropipette was used to isolate and grab the nucleus. A “force” micropipette with a precalibrated deflection spring constant grabbed the nucleus at the opposite end. Using a custom computer program written in LabView, the pull pipette was moved at 50 nm/s to provide extensions of the nucleus up to 6 µm. The computer program tracked the change in distance between the pull and force pipettes to provide the nucleus’ extension (µm) and tracked the deflection of the force micropipette. Using Hooke’s law F = kx where x is the deflection distance and k is the precalibrated bending modulus, the deflection of the force pipette multiplied by the bending modulus (1.2-2 nN/µm) gives the measure of force (nN). Force versus extension was plotted and the slope provided the spring constant (nN/µm) for both the short extension regime (<3 µm) which is dominated by chromatin, and the long-extension regime (>3 µm) which is dominated by lamin-A strain stiffening. Subtracting the short-extension regime spring constant from the long-extension regime spring constant yields the measure of lamin-A based strain-stiffening.

### Imaging and analysis

All images were acquired using Nikon Elements software on a Nikon Instruments Ti2-E microscope, an Orca Fusion Gen III camera, Lumencor Aura III light engine, TMC Clean Bench air table, and a Plan Fluor 40x air DIC M N2 objective (NA 0.75, working distance 0.66 mm, MRH00401). Live-cell imaging was achieved using an Okolab heat, humidity, and CO2 stage-top incubator (H301) in tandem with the Nikon Perfect Focus System. Images were captured using a 16-bit camera on the 40x air objective.

For the immunofluorescence data and the RNA data, the average fluorescent intensities of 30 nuclei were measured per field of view (FOV) and average background was subtracted from each nucleus in Excel. Analysis was done on three FOVs per condition, per replicate (3). For the blebbing rate data, >100 nuclei were counted per condition and the percentage of nuclei that presented with blebs were calculated for each replicate (3) in Excel. Nuclei were counted as blebbed if a protrusion of ≥1 µm in diameter was present with decreased DNA density, as previously outlined (Stephens *et al*., 2018b; Bunner *et al*., 2024; Pujadas Liwag *et al*., 2025).

### Statistics

Data from experimental conditions were considered significant if a student’s *t* test or one-way ANOVA with a post-hoc Tukey test resulted in a p-value of <0.05 when compared to the control data. P values reported as \**P*<0.05; \*\**P*<0.01; \*\*\**P*<0.001, or ns no significance. **Supplemental Figures** NA

## Acknowledgments

We would like to thank Catherin Chu, Nick Lang, Kelsey Prince and Isabel Berg for their helpful and insightful discussions. We thank Masato T. Kanemaki for kindly sharing the mAID-Clover-POLR2A cell line.

## Funding

This work was supported by NIH NIGMS grant Maximizing Investigators’ Research Award R35GM154928.

## Data availability

All figure data can be found posted in the public repository FigShare: https://doi.org/10.6084/m9.figshare.31031863

## Competing interest

The authors declare not competing interests.

## Notes

https://doi.org/10.6084/m9.figshare.31031863

### Competing Interest Statement

The authors have declared no competing interest.

